# Siderophore identification in microorganisms associated with marine sponges by LC-HRMS and a data analytic approach in R

**DOI:** 10.64898/2026.02.09.704990

**Authors:** Alejandro García Ríos, Massuo Kato, Lydia Yamaguchi, Breno Pannia Espósito, Ailan Farid Arenas-Soto

## Abstract

Siderophores are pivotal iron-acquisition biomolecules integral to microbial survival, pathogenicity, and ecology. Elucidating these compounds offers critical insights into the microbial dynamics of marine holobionts and potential therapeutic applications. In this study, we present a culture-independent, data-centric strategy to identify siderophores from the microbiome of three marine sponge species: *Dragmacidon reticulatum, Aplysina fulva*, and *Amphimedon viridis*. Utilizing Liquid Chromatography-High Resolution Mass Spectrometry (LC-HRMS) coupled with a custom R-based analytical workflow (XCMS and MetaboAnnotation), we successfully annotated 59 potential siderophores, 41 of which were confirmed via chromatographic profiling. We employed a rigorous validation pipeline, utilizing multiple iron-adduct calculations [M-2H+Fe]^+^, [M-H+Fe]^2+^, [2M-2H+Fe]^+^, high mass accuracy thresholds (<3 ppm), and retention time precision (CV < 2%). Notably, iron supplementation during extraction did not significantly alter siderophore detection, suggesting constitutive production or environmental saturation. This workflow bypasses the limitations of traditional cultivation, revealing a diverse landscape of iron-chelating metabolites—including Ferricrocin, Aeruginic acid, and Madurastatin—directly within the sponge holobiont.

## Introduction

Siderophores are small, high-affinity iron-chelating compounds produced by microorganisms, including bacteria and fungi. These molecules play a crucial role in microbial survival and pathogenicity by facilitating iron acquisition from the environment. Although iron is an essential nutrient for nearly all living organisms, its bioavailability is limited under aerobic conditions due to its poor solubility. Siderophores overcome this limitation by binding iron with high specificity and affinity, thereby enabling its uptake into microbial cells [1–2].

In a health context, siderophores are of considerable interest due to their roles in both pathogenesis and clinical applications. In many pathogenic bacteria, siderophores act as key virulence factors by enabling efficient iron sequestration in the iron-limited environments of the host, thereby promoting infection and disease progression [3]. Beyond their role in virulence, siderophores exhibit significant therapeutic potential. Their strong iron-chelating properties have been explored for the treatment of iron overload disorders, such as hemochromatosis and certain anemias [4–5]. In addition, synthetic siderophores and siderophore–drug conjugates have been developed to selectively deliver antibiotics into bacterial cells, enhancing treatment efficacy while reducing host toxicity [6].

Siderophores also hold promise for diagnostic applications. Their distinctive biochemical properties can be leveraged to develop sensitive and specific tools for the detection of bacterial infections, particularly those caused by siderophore-producing pathogens [7].

Liquid chromatography coupled with high-resolution mass spectrometry (LC-HRMS) has emerged as a powerful analytical approach for the identification and quantification of siderophores. By integrating the separation efficiency of liquid chromatography with the analytical performance of high-resolution mass spectrometry, LC-HRMS provides a sensitive, specific, and high-throughput platform for detecting siderophores, even within complex biological matrices [8].

Despite LC-HRMS techniques and protocols, some pitfalls still need to be addressed when identifying siderophores. They often occur in low concentrations, especially in complex environments. Their low abundance compared to other metabolites requires sensitive analytical methods for detection. Siderophores have diverse chemical structures that can vary significantly among different organisms. This structural diversity requires comprehensive databases and sophisticated analytical techniques to accurately identify and differentiate siderophores from other molecules [9].

The acquisition, processing, and management of high-resolution spectral data from complex samples are critical for siderophore identification. This process relies on computational tools and statistical methods implemented through established algorithms in specialized software and databases, such as XCMS [10]. These resources support the development of workflows for analyzing high-resolution mass spectrometry datasets, which typically comprise gigabytes of data. The datasets are structured and analyzed using parameters such as retention time, intensity, and m/z, often through freely accessible platforms such as R.

There is a growing demand for fast and cost-effective methodologies capable of detecting siderophore production in complex environmental samples. In parallel, the limited accessibility of curated siderophore databases hampers accurate compound identification [11]. Consequently, accurate siderophore identification depends on efficient computational algorithms capable of processing large datasets and discriminating siderophores from closely related compounds [12].

Symbiotic microorganisms associated with marine sponges, particularly bacteria, represent a rich source of natural products with diverse biological activities, including novel siderophores that remain unexplored in other environments [13]. In this study, we applied an R-based workflow for siderophore detection from high-resolution mass spectrometry data obtained from microorganisms associated with the marine sponges *Dragmacidon reticulatum, Aplysina fulva*, and *Amphimedon viridis*, with the aim of expanding siderophore identification in marine environments. This workflow integrates established R packages, including XCMS, MetaboAnnotation, and ggplot2, for metabolite annotation based on exact mass matching against the current version of the SIDERITE database [14–15].

## Materials and Methods

### Samples

Samples of the sponge species *D. reticulatum, A. fulva*, and *A. viridis* were collected from the São Paulo Atlantic coast (Brazil). We picked 2 grams of the body of each species in triplicate during the summer season (n=36). To investigate inducibility, samples were split into two cohorts: one supplemented with 200 μmol/L^-1^ Iron(III)Ammonium Citrate (FAS) and a control group without supplementation before injection.

### Sample preparation

For sample preparation, 2 g of wet sponge tissue were weighed into 50 mL Falcon tubes and subjected to three successive ethanol extractions (15 mL per extraction), each involving 1 min of vortexing followed by 3 min of sonication at 40 kHz. After centrifugation at 1400 rcf, the supernatants were combined and stored at 4 °C. The resulting 45 mL of ethanolic extract was concentrated using a Buchi Rotavapor R215 with a recirculating chiller (F100) and heating bath (B491) under vacuum (0.074 atm) at 35 °C and 130 rpm, keeping the distillation head below 24 °C. The dried extract was then redissolved in 5 mL of water, thoroughly vortexed, and partitioned with chloroform in four successive 5 mL portions. Each mixture was vortexed for 2 min at 3000 rpm, allowed to equilibrate at 4 °C for 15 min, and centrifuged at 400 rcf for 3 min to separate the aqueous and organic phases. The organic fraction was dried in an Eppendorf SpeedVac Plus (Volatile Heat Mode, no heating) for 5 h, redissolved in 5 mL of methanol, and a 500 µL aliquot was filtered through a pre-conditioned Millipore Ultracel PL-10 centrifugal device at 14000 rcf and 4 °C for 20 min. The filtrate was dried again in the SpeedVac and finally redissolved in 250 µL of dry methanol for LC-HRMS analysis. All steps were performed under controlled temperatures to preserve thermo-labile compounds, providing an efficient and reproducible workflow for the extraction, fractionation, and preparation of sponge-derived metabolites for high-resolution mass spectrometry.

### Analysis by RPLC – HRMS

Liquid chromatography analyses were performed on a Shimadzu LC coupled by electrochemical ionization to an online quadrupole detector with a time-of-flight detector (qToF Bruker Daltonics). The samples of each ultrafiltered organic fraction in methanol (250 µL) were divided into two vials, with 98 µL each. Then, the first vial was added with 2 µL of FAS, 200 µmol L^-1^ (with Fe), and the second, with 2 µL of Milli-Q water (without Fe). An aliquot of 5 µL of the organic fraction with and without iron was injected into column C18 (150 × 3 mm DI, 5 m) Ascentis® express (Sigma-Aldrich) with precolumn. The conditions of the method were water (solvent A) and methanol (solvent B) as mobile phase (both with 0.1% formic acid, v/v) at 0.4 mL min^-1^ and 40°C. The chromatographic conditions were 0 - 10 min (20 to 100%B), 10 - 20 min (100%B), 20 - 23 min (20%B), 23 - 30 min (20%B). All samples (with and without iron addition) were analyzed by LC-qToF with the mass band, m/z from 150 to 2000 Da, at a speed of 1 spectrum s^-1^. The voltage in the capillar was 3500 V; in the fragmenter, 125 V; in the skimmer, 65 V; in the Nozzle 1000 V, and in the octapole, 750 V. Into the ESI, the atomizer conditions were 30 psi at 250°C and 8 L min^-1^ argon.

### Data processing XCMS R Package

36 raw LC-HRMS signals were converted to the standard mzXML format using msConvert from the ProteoWizard tool suite. The converted mzXML files range around 0.3 GB in size per sample. The 36 mzXML files were preprocessed for chromatogram alignment, noise removal, feature detection, and integration. The mass features were extracted using the XCMS 4.0.2 R package standard protocol (https://bioconductor.org/packages/release/bioc/html/xcms.html). We applied 5 centWave peak detection methods, varying the arguments to capture the more specific siderophores in the samples [17] (Supplementary Material S1). The overall centWave peak detection function was defined to find a range between 3 and 60 seconds. Resolution in parts per million (ppm) between 5 and 10, specifying an m/z difference (mzdiff) of 0.001 – 0.005. Further preprocessing involved prefilters with thresholds ranging from 2 to 2000, and noise minimization was capped at 10000. The missing values were placed using the fillChromPeaks function [10, 14]. The final metabolite features table was in a “csv’’ format file.

### Siderophores dataset

Information about siderophores, such as molecular weights and hosted microorganisms, was obtained from the latest version of the SIDERITE database (version 01-20-2025), containing information about 984 unique siderophore structures sourced and curated from the scientific literature [15].

### Adducts Calculated

Previously, we calculated for each “exactmass” from the SIDERITE dataset the adduct [M+H]+ simply by adding the mass of a hydrogen atom. Similarly, an R routine was run to obtain the more plausible iron adducts from each siderophore’s “exactmass”. The iron Adducts were as follows: “[M-2H+Fe]+”, “[M-H+Fe]2+” and [2M-2H+Fe]+.

### Matching Approach with MetaboAnnotation R Package

The mass matching approach consisted of accurately matching the m/z obtained in the metabolite feature table directly against the m/z adducts calculated from the SIDERITE database using the Mass2MzParam and MatchValues functions from the MetaboAnnotation R package 1.6.1 [16], with a predefined tolerance of 0.005 and ppm as low as 3.

### Data Analysis

The information for all m/z matched for siderophores was analyzed using an R code routine for data analysis, including the R library ggplot2.

### Chromatographic peak detection

**Finally, we identified the specific chromatographs matching the m/z detected in the feature table. From the raw mass spectrometry mzXML files** and the m/z obtained in the feature table, we plotted all the chromatogram peaks for all potential siderophores using the function “chromatogram” from the XCMS R package and standard plot drawing functions. This visual inspection of the peaks’ quality works as a criterion to accept as a possible target siderophore. The band width for each m/z was +/- 0.001, and a retention time of 60 seconds was chosen.

## Results

Initially, 62283 peaks were obtained in the feature table after applying the XCMS protocol to the 36 raw mzXML signal files. These m/z were matched against the adducted siderophores in the SIDERITE dataset using the MetaboAnnotation R package. Applying 5 “centWave” functions (by modifying their arguments) yielded 59 potential siderophores (Figures 1 and 2). All the potential siderophores found were adjusted to less than 3 ppm error and more than 5 m/z matches per siderophore (Figures 1 and 2). Figure 1 highlights that different strategies can capture a full metabolomic profile. While Strategy 5 (dark blue bars) was highly effective for detecting compounds like Divanchrobactin and Spoxazomicin C, Strategy 1 (pink bars) was necessary to capture specific isomers of Aeruginaldehyde. This multi-tiered approach yielded 59 distinct potential siderophores, significantly expanding coverage compared to standard single-parameter processing.

**Figure 1.**
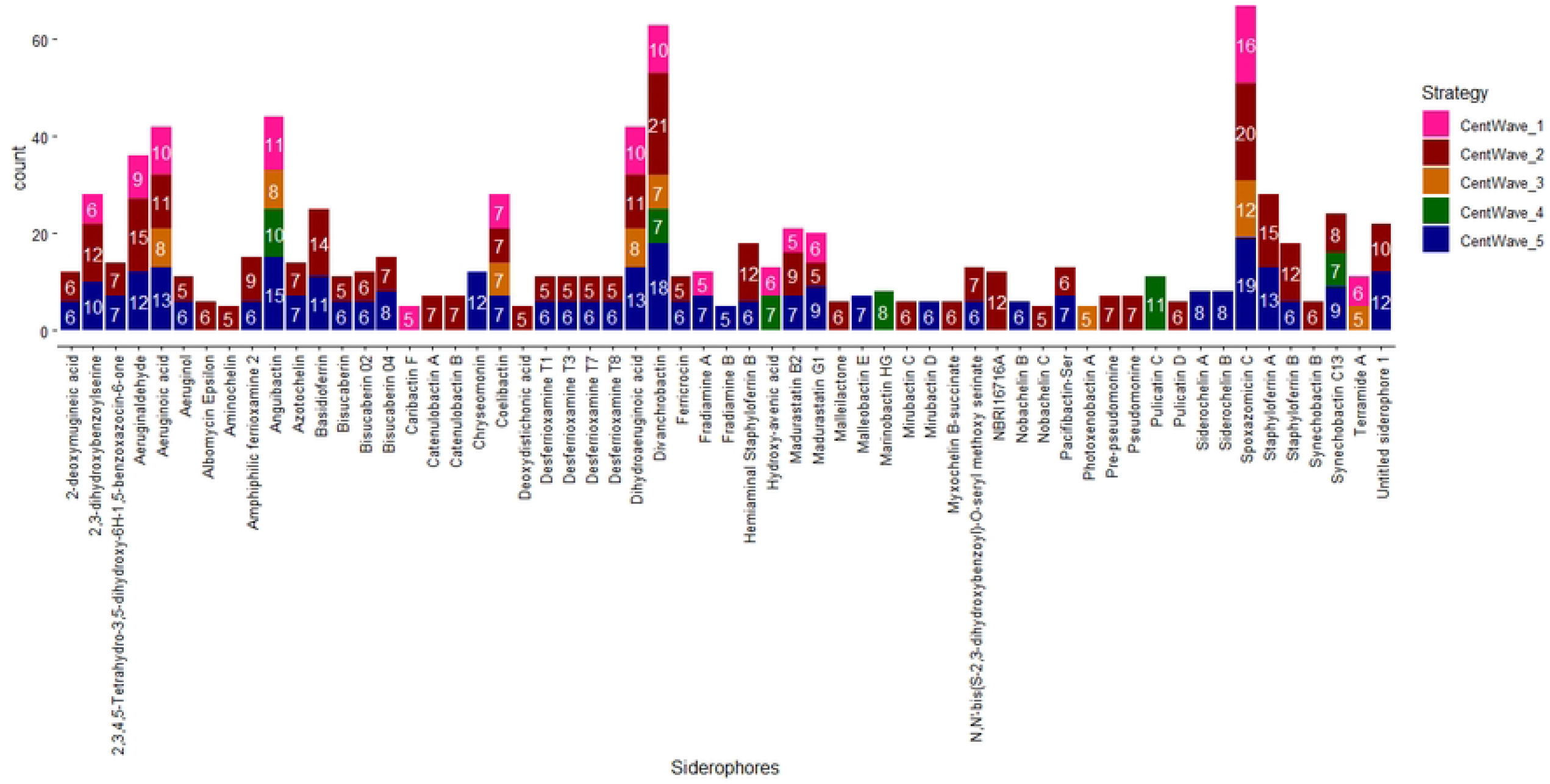
Potential Siderophores matched in the Siderite dataset. Number of potential siderophores found for each centWave function compound detection. It showed only siderophores represented for 5 or more m/z masses per siderophore and with less than 3 ppm error.

**Figure 2.**
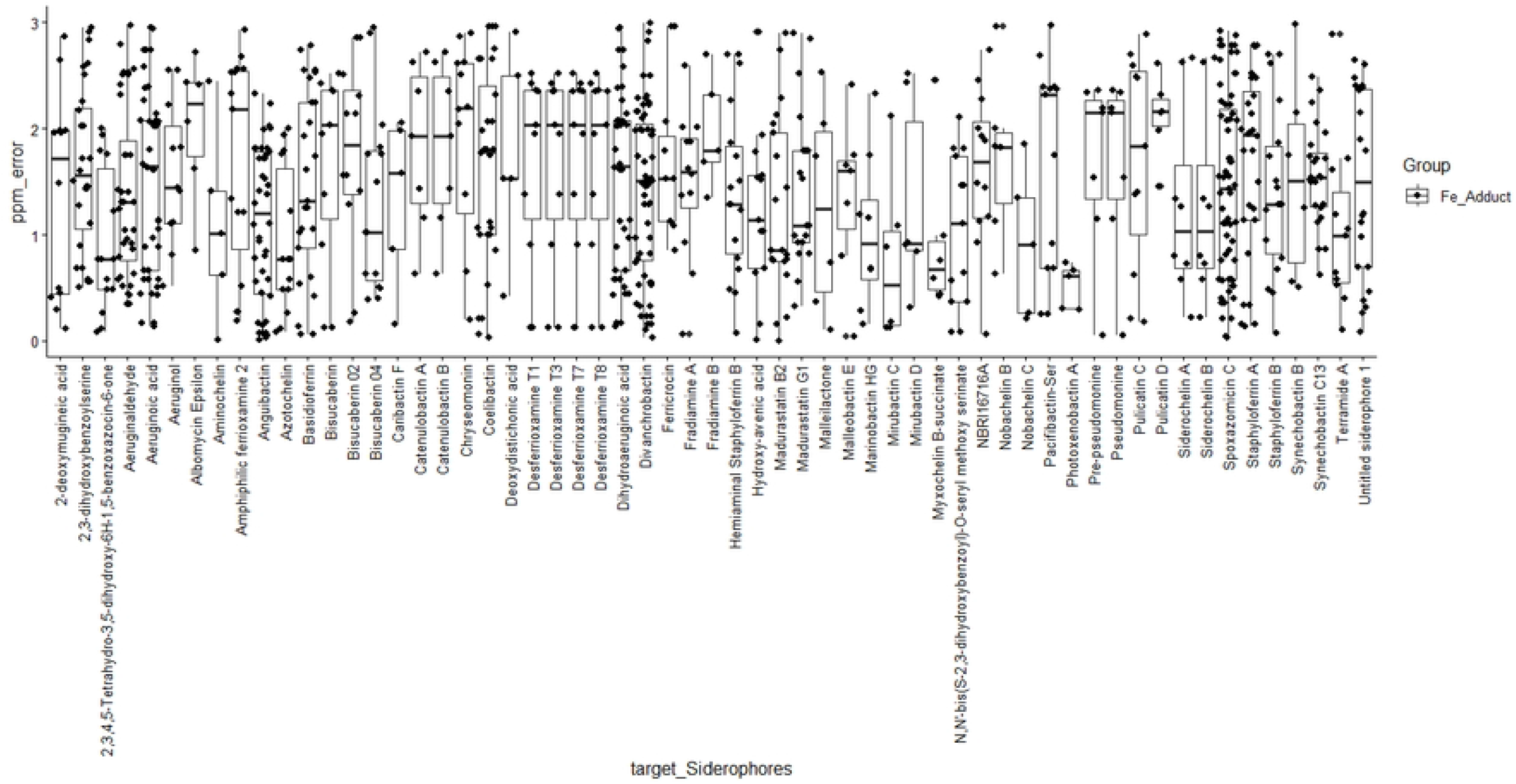
Parts-per-million (ppm) error distribution for each potential Siderophore. Boxplots illustrate the (ppm) error distribution for all 59 annotated potential siderophores.

### Distribution of iron adduct-forming chelating siderophore

We further analyzed the ionization behavior of these compounds. Figure 3 reveals that the singly charged iron adduct [M-2H+Fe]^+^ is the dominant species, accounting for the identification of 31 unique siderophores. This is consistent with the coordination chemistry of hexadentate siderophores, which typically lose two protons to coordinate a ferric ion (Fe^3+^), resulting in a net +1 charge. However, the presence of [M-H+Fe]^2+^ (orange bars) and dimer adducts (pink bars) for compounds like Aeruginic acid indicates diverse ionization pathways that must be accounted for in untargeted screening (Figure 3) [18].

**Figure 3.**
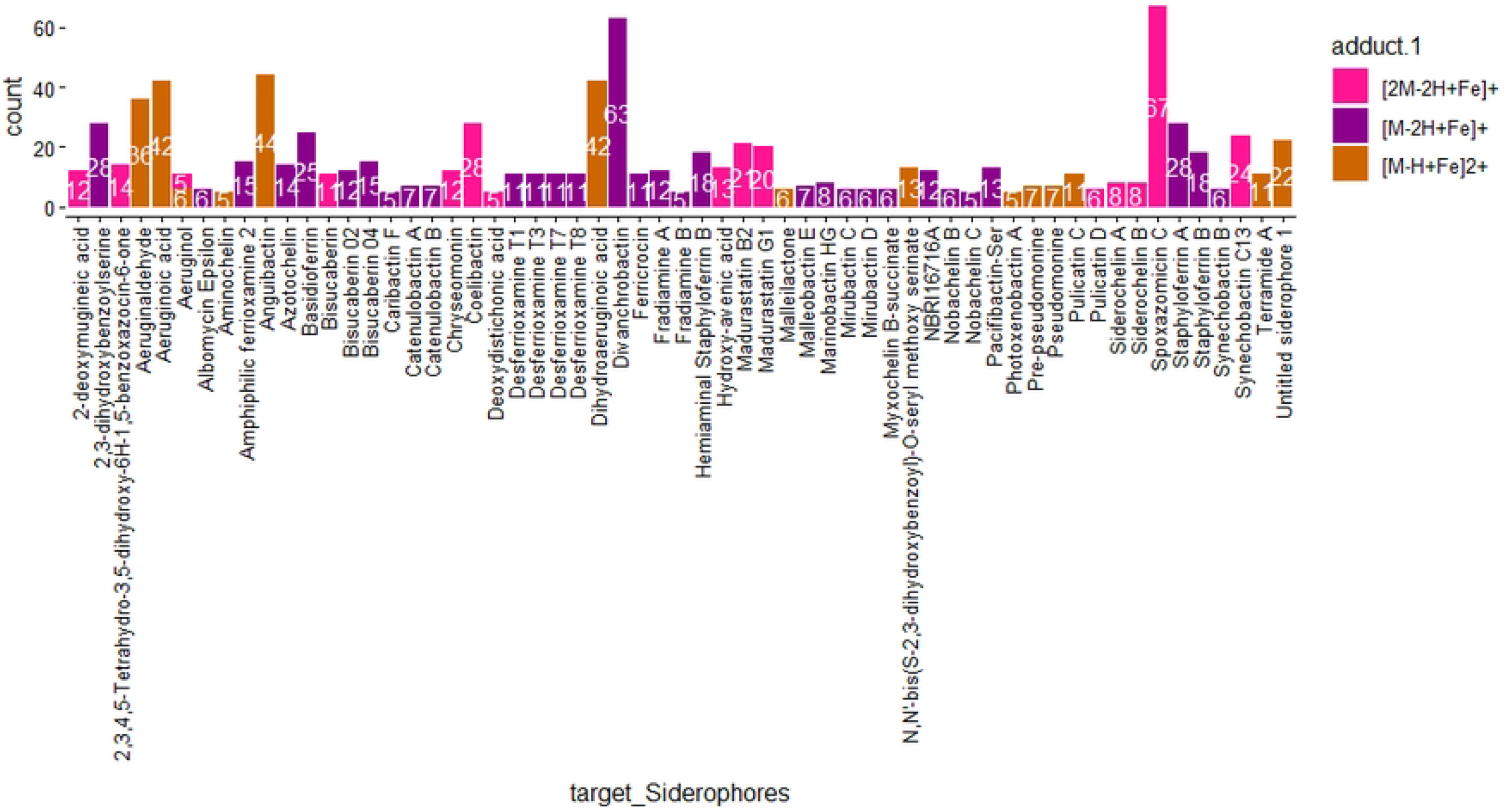
Distribution of (Fe^3+^) ionization adduct-forming. The bar chart illustrates the m/z detected for 59 potential siderophores within (Fe^3+^) ionization adduct-forming. Each bar reflects the frequency of m/z for the siderophores.

### Coefficient of variation (CV) of the retention time for all m/z values of each siderophore

To rule out false positives arising from noise, we assessed retention time (RT) stability as a key criterion for accurately identifying siderophores. The coefficient of variation (CV) provides a standardized measure of RT dispersion for each m/z corresponding to a specific siderophore. Lower CV values indicate tighter clustering of RTs, reflecting greater consistency in the chromatographic behavior of the associated m/z. Our results showed that all 59 siderophores exhibited RT dispersions below 2% (Figure 4), confirming the reliability of their detection.

**Figure 4.**
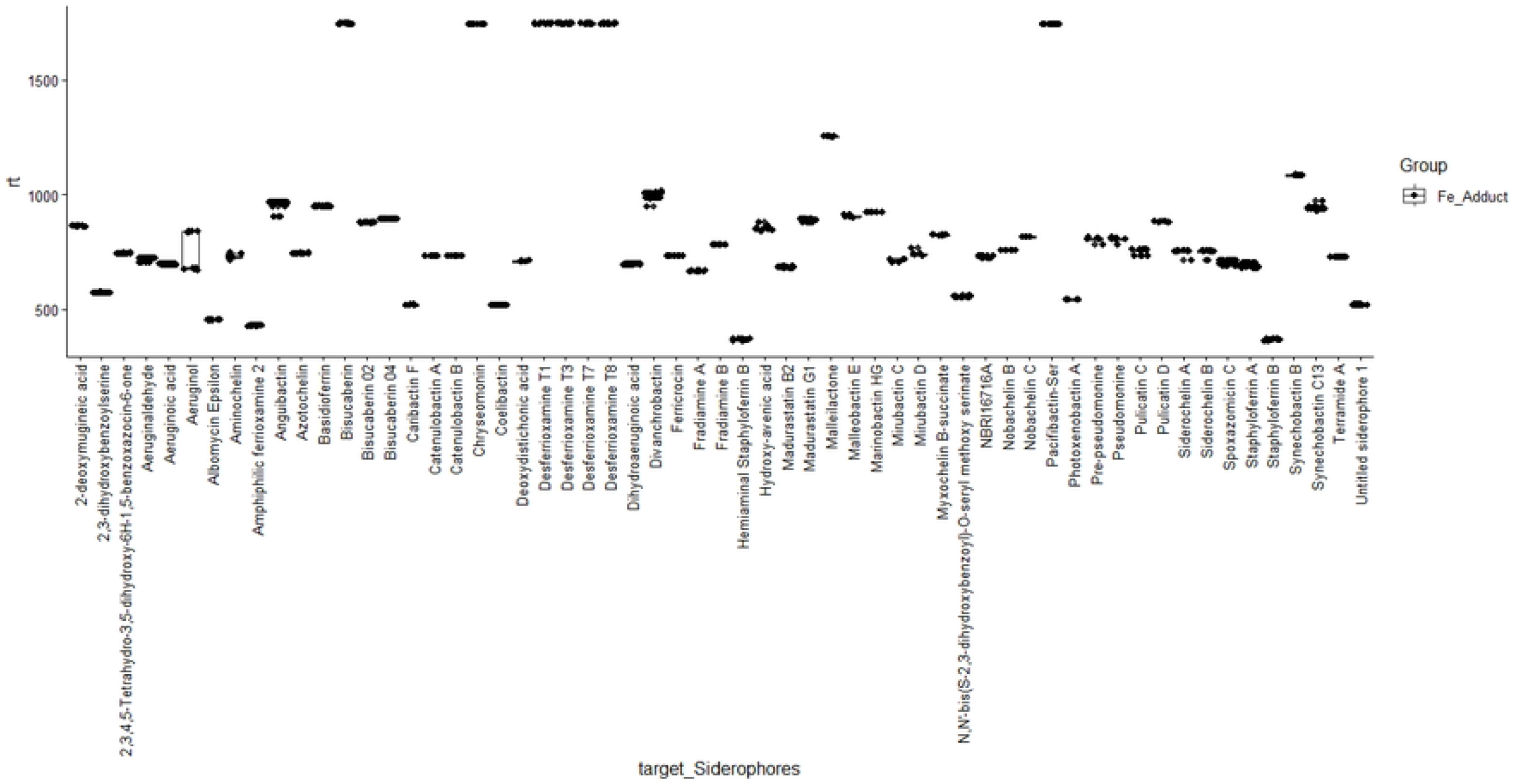
Coefficient of variation (CV) of the retention time (RT) for each m/z corresponding to a specific siderophore. Each short horizontal line represents the RT variation for a given siderophore. Only siderophores with CVs below 2% are shown.

### Effects of iron supplementation on the detection of Siderophore masses

Interestingly, in vitro iron supplementation did not significantly enhance siderophore detection (Figure 5). The counts of detected features were statistically indistinguishable between iron-supplemented (YES) and non-supplemented (NO) samples. This suggests that the identified siderophores in these sponge extracts were already present as iron-complexes (ferri-siderophores) due to the ambient iron in the sponge tissue, or that their production is constitutive rather than strictly inducible under these extraction conditions (Figure 5).

**Figure 5.**
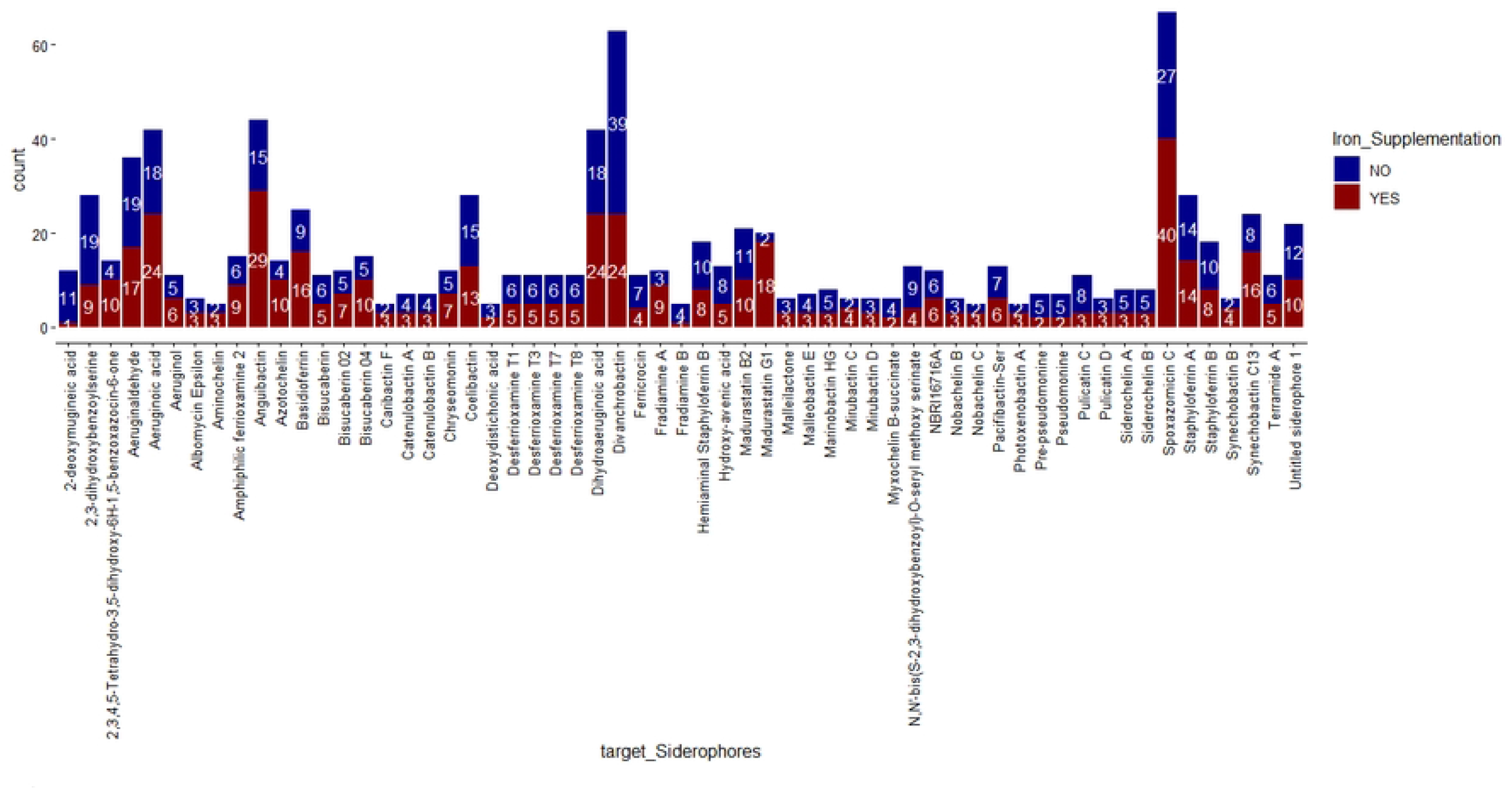
Comparing iron supplementation and non-supplementation on siderophore m/z detected. The bar plot illustrates differences in m/z features for each siderophore when iron was supplemented (“YES”) versus when it was not (“NO”).

### Chromatogram profiles for each detected Siderophore

The definitive validation of our workflow is evidenced by the extracted ion chromatograms (EICs) for each siderophore. Figure 6A and 6B display the elution profiles of the 41 most confident identifications. Most of the chromatograms exhibit sharp, Gaussian peak shapes (e.g., Madurastatin G1 and Nobachelin B), indicative of successful separation from the complex sponge matrix. For many compounds, such as Aeruginaldehyde (Figure 6A), we observe co-elution or slight retention time shifts between the Fe-supplemented (red lines) and non-supplemented (blue lines) traces. This confirms the presence of the apo- and ferri-forms dynamically equilibrating within the column or source. Notably, Figure 6B shows distinct peaks for structural variants, such as the Madurastatin series (B2, G1) and Nobachelin variants. The ability to resolve Madurastatin G1 (RT ∼890s) from Madurastatin B2 (RT ∼680s) demonstrates a well-suited chromatographic method.

**Figure 6.**
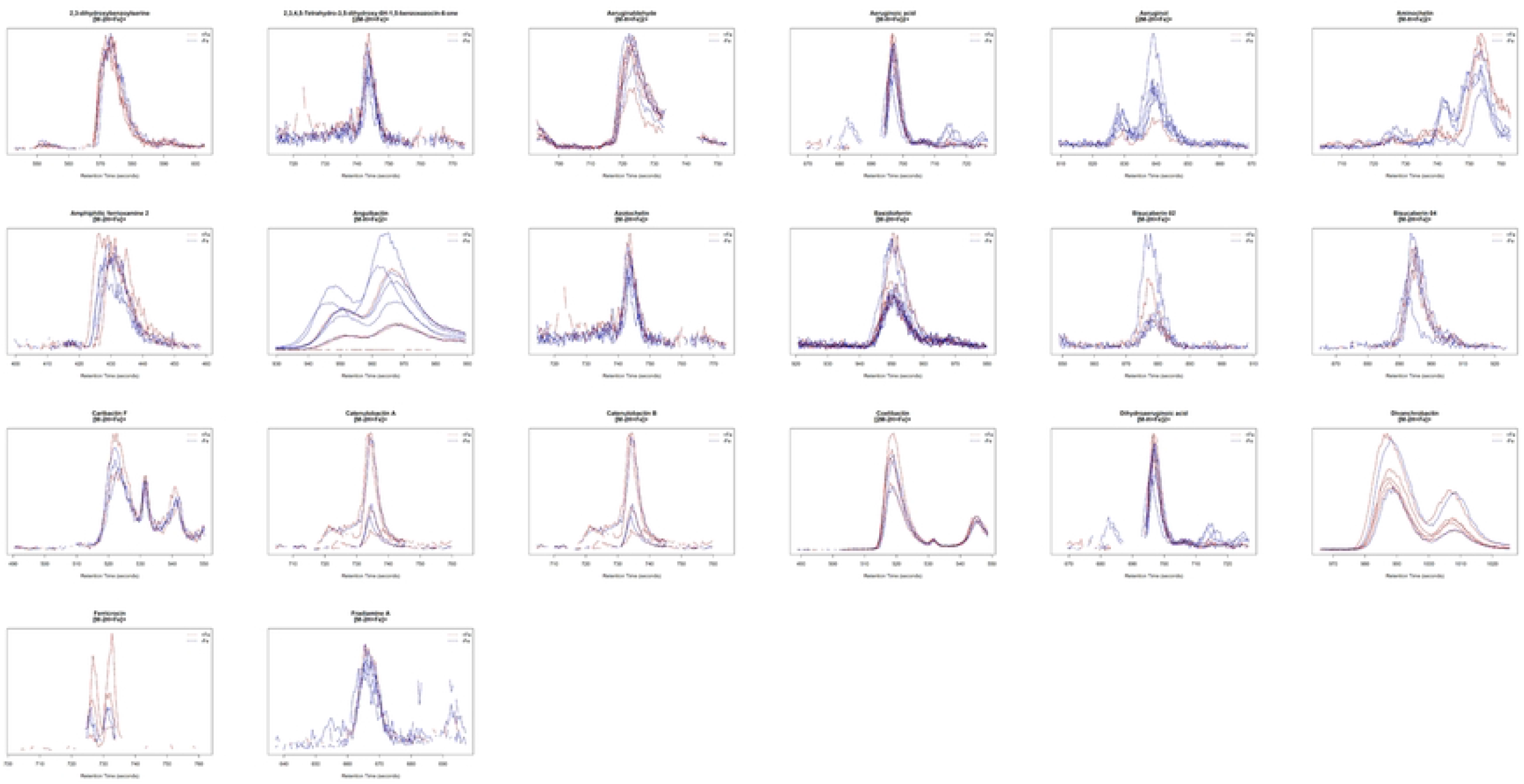
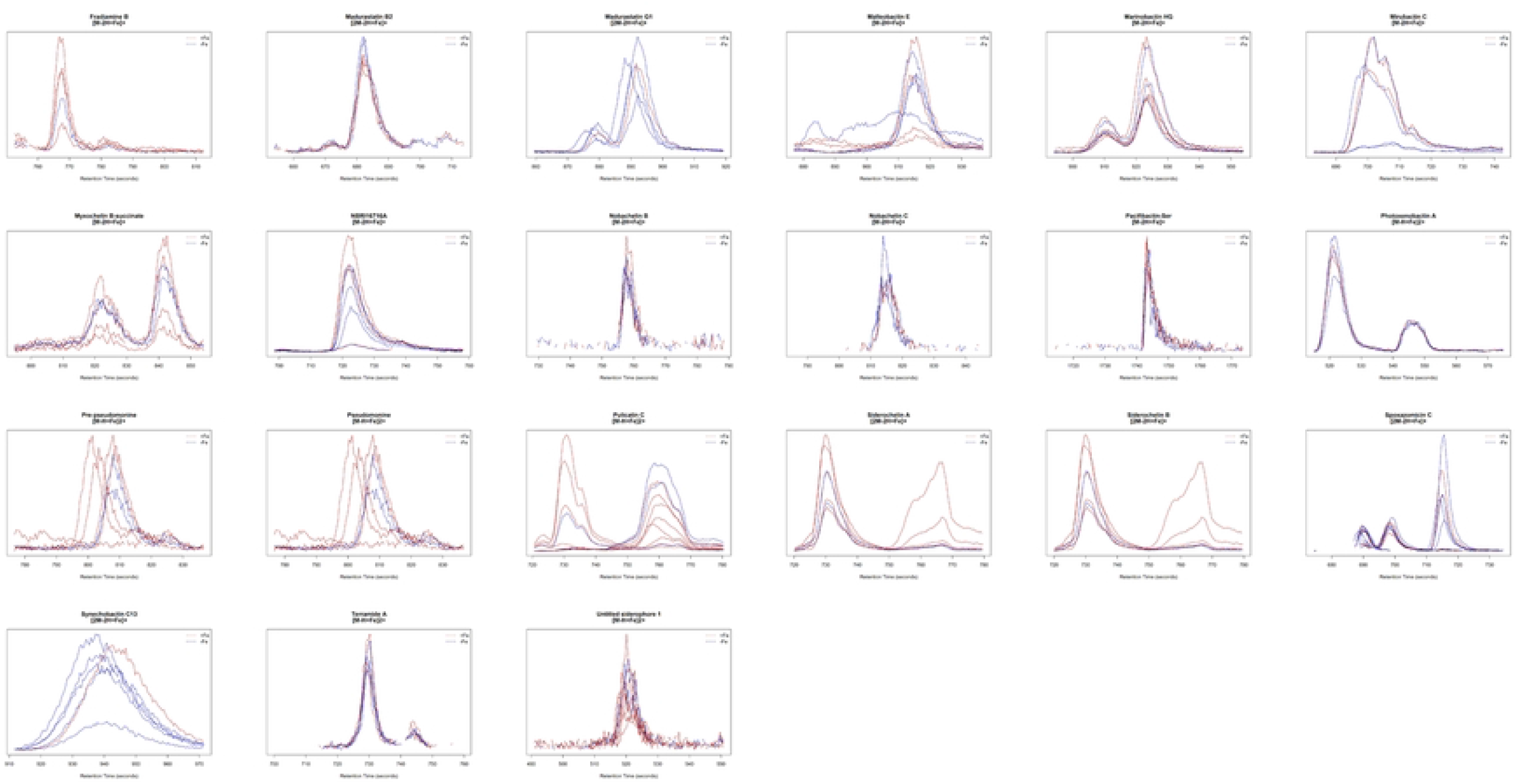
Plots of chromatograms for the more probable siderophores identified in the sponge samples. The bandwidth of all images was +/-0.001 m/z associated with each siderophore, and a retention time of 1 min is shown.

## Discussion

This investigation explores the intricate process of identifying siderophores, which are crucial iron-chelating compounds, derived from microorganisms that, according to this study, are symbiotically associated with marine sponges. We employed a methodology that integrates Liquid Chromatography-High Resolution Mass Spectrometry (LC-HRMS) with data processing and analytical techniques, utilizing specialized R packages such as XCMS [10, 14], MetaboAnnotation [16], and ggplot2. The amalgamation of these analytical tools engenders a thorough and efficient workflow that facilitates the capacity for metabolite discovery from mass spectrometry data. This approach implements a mass-feature matching protocol against a curated and highly specialized siderophore database [15], which resulted in the identification of 41 potential siderophores directly obtained from the biological samples of three distinct sponge species, all accomplished without the necessity for the prior isolation or cultivation of the associated microorganisms (Figures 1 to 5).

We executed a meticulous filtration process grounded in a set of diverse criteria delineated in our approach. These specified criteria include the calculation of three distinct iron adducts, a parts per million (ppm) resolution threshold of less than 3 for all feature adducts, m/z feature counts per siderophore greater than 5, a coefficient of variation in the retention time that does not exceed 2% across all m/z features associated with each identified siderophore (Figure 2 to 4), and the plotting of the chromatograms (Figure 6a and 6b). Implementing these stringent criteria significantly enhances the reliability of our identification process for these specialized chemical compounds (Supplementary Material S2).

Although there exists a plethora of tools specifically designed for the identification of metabolites derived from liquid chromatography-mass spectrometry (LC-MS) experiments, which includes notable programs such as MetaboAnalyst [19], MS-DIAL [20], MZmine [21], OpenMS [22], MetFrag [23], GNPS [24], and MetaboLights [25], it is important to recognize that programmatic approaches that are implemented using programming languages like R still possess distinct advantages, particularly in terms of their inherent flexibility and adaptability to various analytical scenarios. Furthermore, the utilization of specialized packages such as ggplot2 significantly enhances the visualization capabilities of data, thereby facilitating the creation of highly customized graphics that are specifically tailored to meet specific analytical requirements, ultimately leading to improved interpretability and more effective presentation of the resultant information derived from complex datasets (Figures 1 to 5).

There is still room for exploration of siderophores within various environmental and biological matrices. Towards this objective, earlier studies involved the isolation and cultivation of microorganisms before the execution of LC-MS experiments. For instance, Mawji et al. 2010 discerned 22 siderophores derived from the Atlantic Oceanic waters, enriched with various compounds serving as carbon and nitrogen substrates, thereby enhancing the capture of siderophores [26]. Lehner et al. 2013 conducted a comprehensive investigation that led to the identification and subsequent detection of 18 distinct iron chelators produced as a result of the laboratory cultivation of a collection comprising 10 wild-type strains of the *Trichoderma* genus [27]. Baars and colleagues, in their research conducted in 2015, identified a total of 35 distinct siderophores that were produced by the cultivation of the nitrogen-fixing soil bacterium *Azotobacter vinelandii*, which is widely recognized for its significant role in the nitrogen cycle [28]. Boiteau et al. 2019 identified 9 siderophores in culturing microorganisms from soil samples and reported the siderophore Schizokinen, also found in our screening [29]. Finally, a publication released in the year 2022 by Liu and coauthors pursuing a similar research objective reported 10 Siderophores isolated from the actinomycete *Streptomyces diastaticus* NBU2966, which is associated with one marine sponge belonging to the Axinellida order [30]. Our study represents one of the scarce instances in the scientific literature investigating the presence of siderophores linked to microorganisms that inhabit the ecological niche provided by sponges [20]. Our study contribution to this field significantly enhances the existing body of knowledge about the presence and relevance of siderophores in marine environments that are specifically associated with species of sponges.

Marine sponges are known to harbor diverse microbial communities, including bacteria and fungi, which play crucial roles in the sponge holobiont. The literature concerning the identification of siderophores synthesized by sponge-associated microbial communities is quite sparse. A particular investigation revealed that microorganisms extracted from sponges exhibited no detectable production of siderophores, even when iron was supplemented to the culture medium [31]. Nevertheless, certain isolated bacterial strains demonstrated the capability to produce siderophores when stimulated by exogenous siderophores, suggesting a potential signaling function of exogenous siderophores in enhancing the production of indigenous siderophores in marine bacterial populations [31]. Concerning our experimental methodology, we noted that *in vitro* stimulation with iron did not result in a significant difference in the count of m/z features of siderophores subjected to iron stimulation (Figure 5). We suspect that most of the identified siderophores are strong iron-chelator compounds, and probably they were already forming chelating complexes, and no increasing chelation activity was observed with exogenous iron supplementation (Figure 5).

Based on the 41 distinct siderophores identified in this investigation, we postulate that sponges represent promising reservoirs for siderophores within their native ecosystems, particularly given the prevalent iron scarcity in seawater and the intense competition for this vital metal. The symbiotic interaction between sponges and their associated microorganisms may be fundamentally influenced by the distinctive ecological niches present within the sponge species [32].

## Conclusions

Our study employed a homemade, data-driven approach combining Liquid Chromatography-High Resolution Mass Spectrometry (LC-HRMS) with data processing in R using packages such as XCMS, MetaboAnnotation, and ggplot2 to identify siderophores in marine sponge samples. This methodology detected 41 distinct siderophores across three sponge species: *Dragmacidon reticulatum, Aplysina fulva*, and *Amphimedon viridis*, without the need for prior isolation or cultivation of associated microorganisms. This research highlights the potential of marine sponges as reservoirs for diverse siderophores, which could serve as indicators of marine environmental quality and ecosystem health. *In vitro* iron stimulation did not significantly alter siderophore m/z values. This research contributes to the limited literature on siderophores associated with marine sponges.

## Acknowledgments

This work was supported by Universidad del Quindío (100016837) under Grant Res. No.048 (19 July 2017); the funding agencies CAPES, for the PAIS scholarship, process 88882,328227/2019-01 for study expenses and technical reserves; FAPESP for research financing; to Dr. Márcio Reis Custódio (Instituto de Biología, USP); CEBIMAR for the samples and to Dr. Pedro Vidinha (Laboratório de Química Ambiental, IQ-USP).

## Supplementary material

Supplementary Material S1: Contains the centWave R functions with the arguments used in this study

Supplementary Material S2: Contains the R scripts for the basic siderophore identification.

